# Hippocampal Commissural Circuitry Shows Asymmetric cAMP-Dependent Synaptic Plasticity

**DOI:** 10.1101/2025.06.11.659103

**Authors:** Lukas Faiss, Aikaterini Salivara, Silvia Oldani, Jörg Breustedt, Dietmar Schmitz, Benjamin R. Rost

## Abstract

Hemispheric asymmetries in NMDAR-dependent synaptic plasticity have been described in hippocampal area CA1, but it remains unclear whether similar lateralized mechanisms exist for cyclic adenosine monophosphate (cAMP)-dependent plasticity. Here, we investigated whether cAMP-mediated potentiation of synaptic transmission in mouse CA1 exhibits hemisphere-specific properties. In recordings with electrical stimulation of CA1 inputs, a subset of recordings in the left, but not in the right hemisphere CA1, exhibited a pronounced cAMP-induced potentiation of field excitatory postsynaptic potentials (fEPSPs). To isolate input specific contributions, we expressed the optogenetic actuator ChrimsonR unilaterally in the CA3/CA2 region of wild-type mice. Light-evoked glutamate release from ipsilateral Schaffer collaterals showed no cAMP sensitivity in either hemisphere, while commissures originating from the right (COR) exhibited cAMP-mediated potentiation of transmission in a subset of experiments. Notably, this effect was absent at commissures originating from the left (COL). The selective presence of the effect prompted us to further investigate the underlying cell population using CA3-specific (G32-4 Cre) and CA2-specific (Amigo2-Cre) driver lines. Recordings from synapses of CA3 COR recapitulated the cAMP-induced potentiation of transmitter release observed in wild-type animals. However, the effect was again restricted to a subset of experiments and was absent in recordings with specific stimulation of CA2 COR. Our results demonstrate a variable cAMP-sensitivity of transmitter release at COR synapses in the left CA1. Altogether, we reveal a hemisphere-specific cAMP-mediated synaptic plasticity at CA3 COR onto CA1, underscoring hidden heterogeneity and lateralization in hippocampal circuit function.

## Introduction

Within the hippocampus, CA1 pyramidal neurons receive glutamatergic input from both ipsilateral Schaffer collaterals and contralateral commissural projections, both originating from CA3 and CA2 pyramidal neurons (Blackstad, 1956; Andersen, Blackstad and Lömo, 1966; Gottlieb and Cowan, 1973). As Schaffer collaterals and commissural fibers share similar targets in the CA1 stratum radiatum, their overlapping distributions complicate selective stimulation (Swanson, Wyss and Cowan, 1978). Despite the mirror-image wiring at the macroscopic level, CA1 inputs appear to exhibit lateralized, hemispheric-specific functional differences. In mice, CA1 pyramidal cell synapses show input specific expression of ionotropic glutamate receptors. Synapses with presynaptic terminals originating from the right hippocampus predominantly express GluA1-containing AMPA receptors, whereas synapses with input from the left hippocampus show higher expression levels of GluN2B-containing NMDA receptors (Kawakami *et al*., 2003; Shinohara *et al*., 2008). In *inversus viscerum* (iv) mice, hemispheric asymmetry is absent. This lack of lateralization is linked to deficits in learning and memory, including slower spatial learning and reduced working memory performance (Kawakami *et al*., 2008; Goto *et al*., 2010). Gene expression analyses revealed greater transcriptional changes in the right hippocampus following spatial learning (Klur *et al*., 2009). Additionally, disrupting interhemispheric connections uncouples gamma wave synchronization, suggesting that the right hippocampus integrates bilateral information before cortical transmission (Benito *et al*., 2016). Hemispheric asymmetries also influence synaptic plasticity. NMDAR-dependent long-term potentiation (LTP) at CA1 synapses was suggested to be preferentially induced by left CA3 inputs, implicating a dominant role of the left hemisphere in driving postsynaptic LTP (Kohl *et al*., 2011; Shipton *et al*., 2014; El-Gaby, Shipton and Paulsen, 2015). Furthermore, commissural projections from CA2/3 in the left hemisphere are essential for full postsynaptic plasticity and place field formation in CA1 (Fan *et al*., 2023). While some findings support this left-dominant model, its generality across species and conditions remains debated (Martin, Shires and da Silva, 2019; Kuwajima *et al*., 2020). Beside the classical NMDA receptor/CaMKII-dependent signaling cascade, cAMP-dependent pathways represent additional mechanisms for inducing or supporting both pre- and postsynaptic forms of plasticity (Bernabeu *et al*., 1997; Otmakhova *et al*., 2000; Otmakhov *et al*., 2004). Here, we investigated whether Schaffer collateral or commissural CA3-CA1 synapses exhibit hemispheric specific cAMP-dependent potentiation of synaptic transmission. Using optogenetics and electrophysiological recordings, we tested whether pharmacological cAMP modulation differentially affects synaptic plasticity in CA1.

## Results

### Asymmetrical cAMP modulation of hippocampal CA1 inputs

Pharmacological enhancement of cAMP signaling has previously been reported to increase synaptic input to CA1 pyramidal neurons by approximately 20% (Muñoz and Solís, 2019). To assess whether this modulation differs across hemispheres or specific input pathways, we defined a >20% increase in synaptic transmission following bath application of 50 µM forskolin (FSK), a pharmacological activator of adenylyl cyclases, as a threshold for a positive cAMP effect. Using this criterion, we examined hemisphere-specific responses by recording field excitatory postsynaptic potentials (fEPSPs) in area CA1 of both the left and right hippocampus (Fig. 1A, B). In the left hippocampus, a subset of recordings exhibited potentiation (>20% increase), whereas recordings from the right hemisphere consistently failed to meet this threshold. Specifically, 7 out of 29 recordings in the left hemisphere (28.6%) showed a significant FSK-induced increase in synaptic transmission (mean effect: 138.3 ± 6.9%, n = 7), while the remaining recordings showed no effect (99.5 ± 2%, n = 22)(Fig. 1C, F). In contrast, none of the 28 recordings from the right hemisphere surpassed the 20% threshold (overall mean: 99.6 ± 9.5%, n = 28) (Fig. 1D, F). Statistical comparison confirmed a significant difference in FSK responses between hemispheres (p = 0.018, unpaired t-test with Welch’s correction, Fig. 1E). To further quantify this effect, we used a bootstrap approach to calculate Cohen’s d, yielding a mean value of 0.664, indicative of a medium effect size and supporting the distinction between the two hemispheric datasets (Fig. 1G). Together, these results reveal a clear variability in cAMP responsiveness, with FSK enhancing synaptic transmission only in a subset of left hemisphere recordings. The absence of comparable effects in the right hemisphere suggests a lateralized expression of cAMP-dependent plasticity. Electrical stimulation activates both Schaffer collateral and commissural inputs to CA1, raising the question of whether the observed potentiation is specifically mediated by one of these pathways.

**Figure 1.**
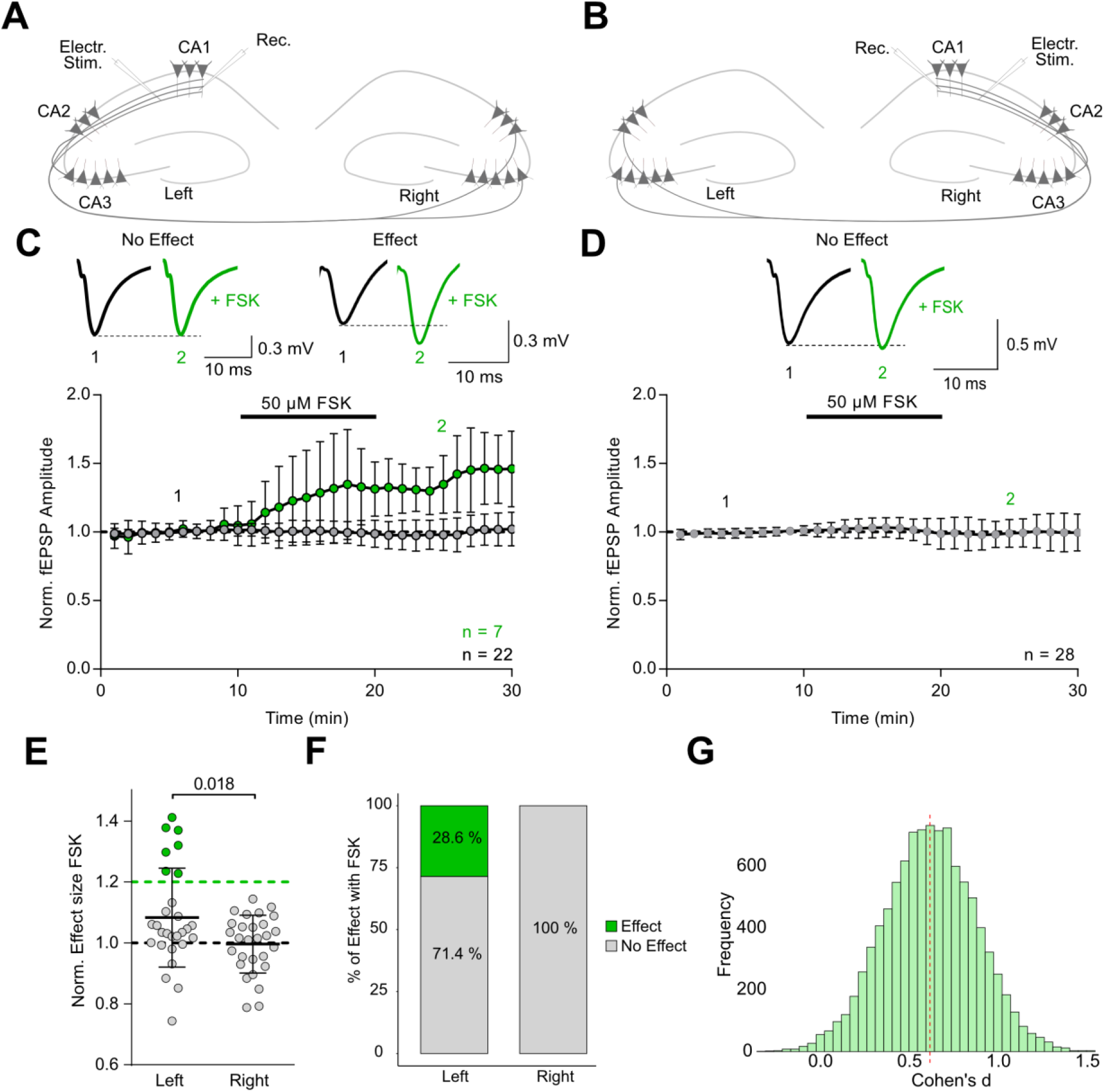
Selective potentiation of synaptic responses by forskolin in the left hippocampus. **A, B** Schematic of the field stimulation and recording setup in the CA1 region of the left **(A)** and right **(B)** hippocampus. **C, D** Representative traces and time courses of electrically evoked fEPSPs recorded in the left **(C)** and right **(D)** hippocampus. Black traces represent average of a 10-min baseline; green traces show responses after application of 50 μM forskolin (FSK). The averages of the recordings with an FSK-effect >20% are shown in green, while the averages of all other recordings are shown in grey (mean ± SD). **E** Summary of FSK effects on normalized fEPSP amplitudes reveals significantly greater potentiation in the left hemisphere compared to the right (left: 108.3% ± 16.3%; right: 99.6% ± 9.5%; *p* = 0.018, unpaired t-test with Welch’s correction). **F** Percent stacked bar plot indicating the distribution of recordings with and without significant potentiation, based on a 20% threshold (Left: 28.6% of recordings showed an effect, 71.4% no effect; Right: 0% with effect). **G** Histogram of bootstrapped Cohen’s d values comparing left and right hemisphere recordings, revealing a medium effect size favoring the left hemisphere (Cohen’s d = 0.664).

### Selective Activation of Commissural Fibers Reveals Asymmetric cAMP Modulation

To dissect both hemisphere- and pathway-specific effects of cAMP signaling, we employed optogenetic activation of Schaffer collateral and commissural inputs using the channelrhodopsin ChrimsonR (K176R) (Klapoetke *et al*., 2014). An AAV9 vector encoding hsyn-ChrimsonR-tdTomato was unilaterally injected into either the left (Fig. 2A, B) or right (Fig. 2B) CA3/CA2 region of wild type (WT) mice. Three to five weeks post-injection, acute hippocampal slices were prepared, and ChrimsonR expression was verified via tdTomato fluorescence (Fig. 2C, D). Synaptic responses were recorded in CA1 stratum radiatum, either ipsilaterally or contralaterally to the injection site, using brief 590-nm light pulses (0.5–2 ms, 2–4 mW/mm^2^) delivered via the objective near the recording electrode at 0.1 Hz. We first evaluated the effect of cAMP elevation on Schaffer collateral synapses. Bath application of FSK failed to induce consistent potentiation in either hemisphere. In the left hippocampus, none of the recordings (0 out of 9) surpassed the 20% threshold (mean response: 103.0 ± 7.1%, n = 9, N = 4, Fig. 2E, H). Similarly, in the right hemisphere, 19 of 20 recordings showed no effect (mean: 99.9 ± 9.9%), while one recording (5.3%) exhibited an increase of synaptic transmission above the threshold (124.5 ± 8.4%) (Fig. 2F, H). Although a single right hemisphere recording showed a >20% increase, statistical analysis revealed no significant difference in overall FSK responses between both hemispheres (p = 0.415, unpaired t-test, Fig. 2G). A bootstrap analysis of effect size yielded a Cohen’s d of 0.354, indicating a small effect and reflecting the marginal difference between groups (Fig. 2I). These findings suggest that Schaffer collateral pathways generally lack robust cAMP sensitivity in both hemispheres.

**Figure 2.**
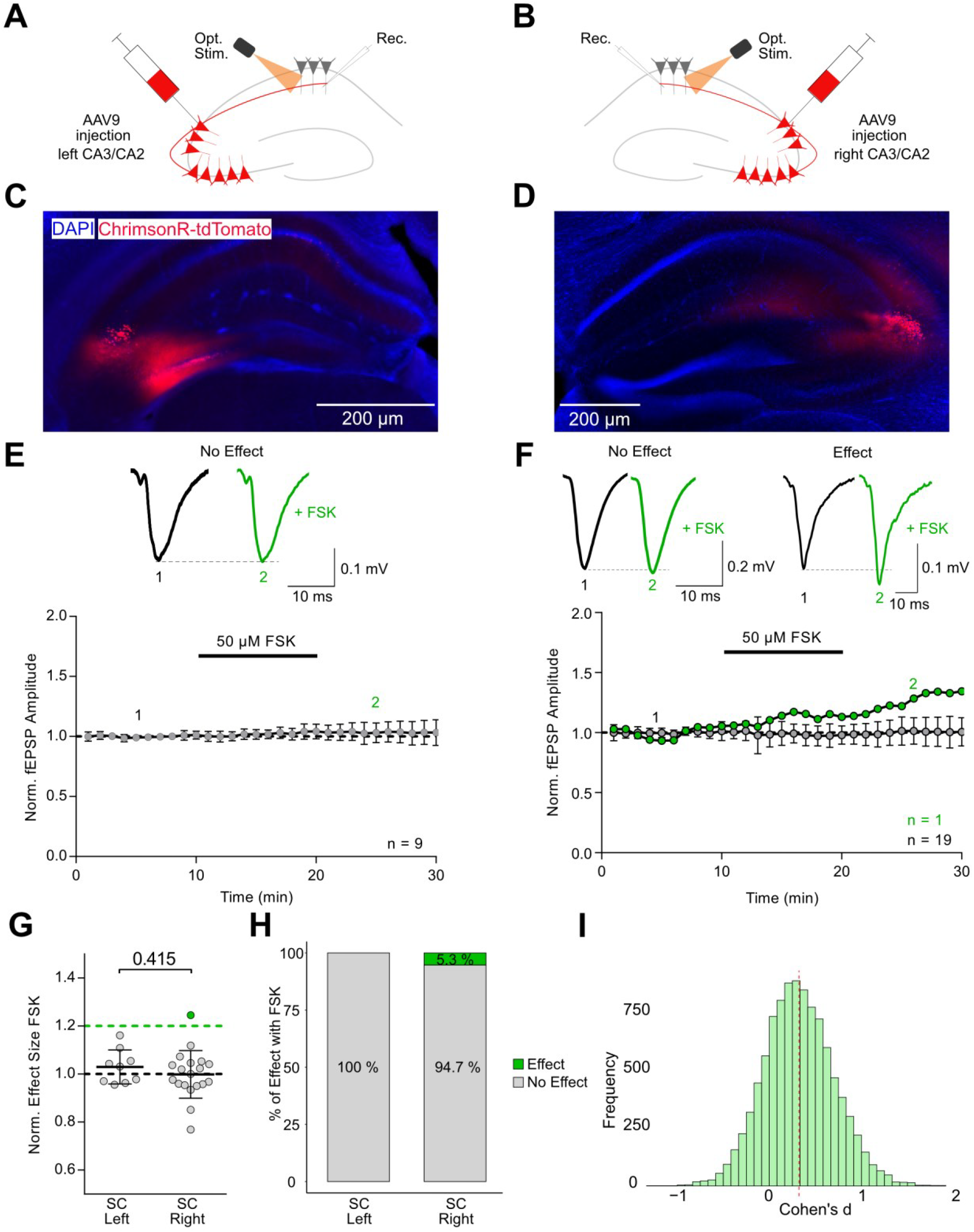
No FSK effect on Schaffer collateral transmission evoked by optogenetic stimulation. **A, B** Experimental configuration for selective optogenetic stimulation of Schaffer collateral inputs with recordings from the CA1 region in the left **(A)** or right **(B)** hippocampus. **C, D** Fluorescent micrographs showing ChrimsonR-tdTomato expression in the CA3/CA2 region in the left **(C)** or right **(D)** hippocampus. DAPI staining labels cell nuclei. **E, F** Representative optogenetically-evoked fEPSP traces from the left **(E)** and right **(F)** hippocampus. Black traces represent baseline responses; green traces show post-FSK responses. Time courses from the left **(E)** and right **(F)** hippocampus. Time course plots distinguish effect recordings (>20% increase, green) from non-effect recordings (grey). **G** Group summary of normalized fEPSP amplitudes shows no significant difference in FSK-induced potentiation between hemispheres (*p* = 0.415, unpaired t-test; SC = Schaffer Collaterals). **H** Percent stacked bar plot categorizing individual recordings based on presence or absence of a >20% increase in synaptic strength. No recordings from the left hemisphere met this criterion, while 1 out of 20 recordings (5.3%) in the right hemisphere showed an effect. **I** Distribution of bootstrapped Cohen’s d values comparing left and right hemisphere responses indicates a small effect size (Cohen’s d = 0.354), indicating a minor hemispheric difference.

In contrast, optogenetic activation of commissural fibers revealed substantial variability in cAMP responsiveness, particularly depending on the hemisphere of origin. At commissures originating from the left (COL) and terminating at the right hemisphere CA1, FSK application failed to elicit potentiation exceeding the 20% threshold in any of our recordings (mean response: 103.2 ± 6.4%, *n* = 20, *N* = 5; Fig. 3E). This lack of effect was consistent across all recordings, indicating minimal variability within this group. Conversely, commissures originating from the right (COR) displayed much greater heterogeneity. While the group mean was only modestly elevated (114.0 ± 23.4%, *n* = 28, *N* = 11; Fig. 3G), this was driven by a subset of recordings that showed strong potentiation. Specifically, 6 out of 28 recordings from COR (21.4%) exceeded the 20% effect threshold (mean: 155.6 ± 6.3%). The remaining 22 recordings (78.6%) exhibited no change (mean: 107.0 ± 2.8%), highlighting a substantial inter-recording variability (Fig. 3F). A statistical comparison of overall responses revealed a trend toward greater cAMP sensitivity in COR projections (*p* = 0.085, unpaired t-test; Fig. 3G), although this did not reach conventional significance. To better capture the effect despite this variability, we conducted a bootstrap analysis, which yielded a Cohen’s d of –0.591, indicative of a medium effect size and suggesting a functional asymmetry between the commissural fibers of both hemispheres (Fig. 3I). Notably, the presence of both strongly responsive and non-responsive recordings within the COR suggests that not all COR projections are sensitive to cAMP. This variability likely reflects differences in the underlying cellular sources of these projections. Since AAV-driven ChrimsonR-tdTomato expression in WT mice lacks specificity for distinct pyramidal subtypes, the observed variability raises the possibility that the subset of cAMP-sensitive commissural fibers originate from a specific group of pyramidal neurons in the right hemisphere. Determining the identity of these responsive cells will be essential for understanding the basis of this asymmetric and heterogeneous plasticity.

**Figure 3.**
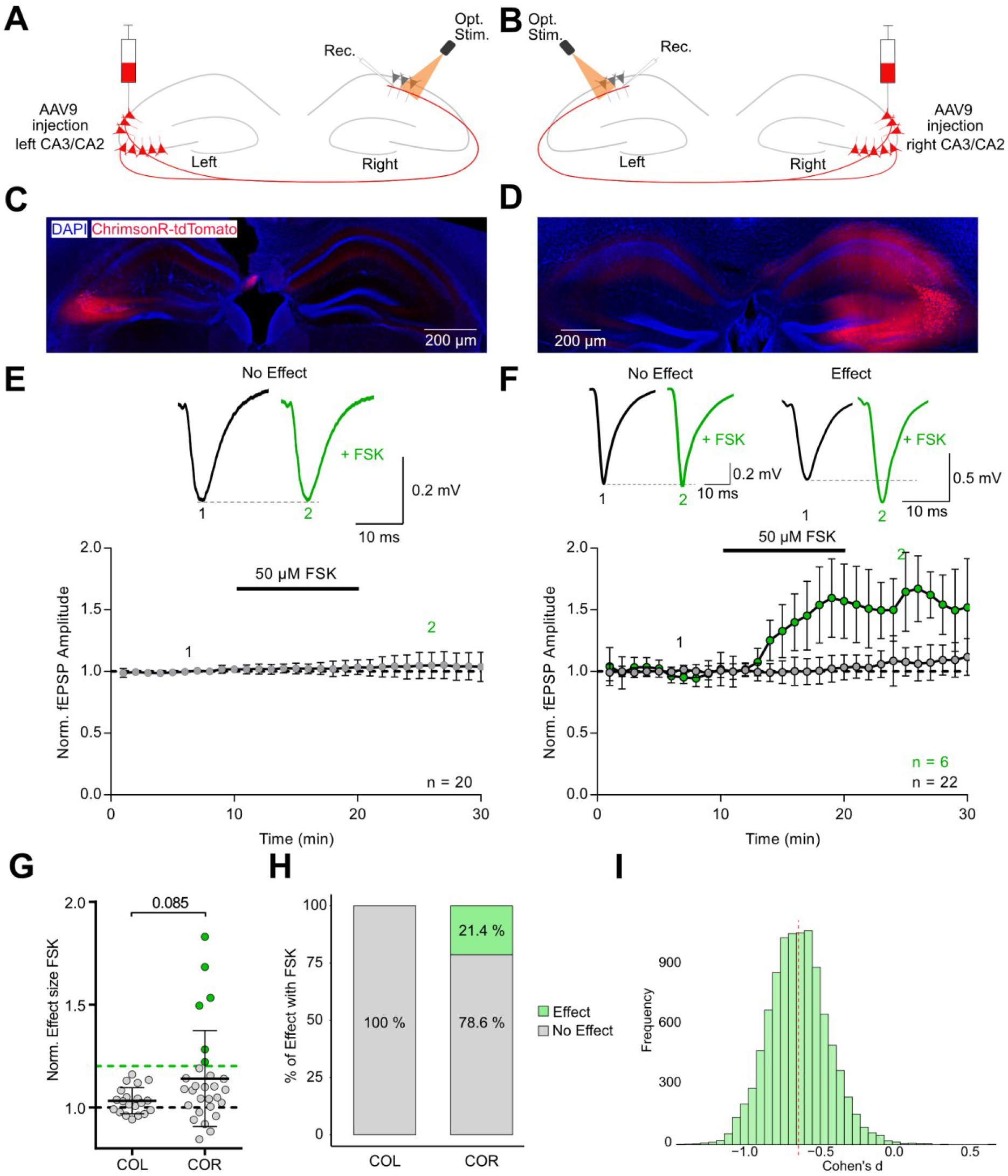
cAMP selectively enhances transmission at synapses formed by commissures originating from the right. **A, B** Optogenetic setup for stimulating commissural fibers originating from the left **(COL, A)** or right **(COR, B)** hippocampus. **C, D** Fluorescent images of ChrimsonR-tdTomato expression in CA3/CA2 pyramids of the left **(C)** and right **(D)** hemisphere. **E, F** Representative traces and timeplot of fEPSPs evoked by optogenetic stimulation of synapses formed by COL **(E)** and COR **(F)**. Black traces represent baseline responses; green traces show post-FSK responses. Time course plots distinguish effect recordings (>20% increase, green) from non-effect recordings (grey). **G** Group summary of normalized fEPSP amplitudes illustrates a selective occurrence of FSK-induced potentiation of transmission at synapses formed by COR (*p* = 0.085, Mann-Whitney testt). **H** Percent stacked bar plot categorizing individual recordings based on presence or absence of a >20% increase in synaptic strength. No recordings from the left hemisphere met this criterion, while 6 out of 28 recordings (21.4%) in the right hemisphere showed a pronounced cAMP-effect. **I** Distribution of bootstrapped Cohen’s d values comparing left and right hemisphere responses indicates a medium effect size (Cohen’s d = -0.591), suggesting a hemispheric difference.

### CA3, but Not CA2, Commissural Projections Exhibit cAMP-Dependent Potentiation

To further pinpoint the origin of commissural fibers underlying FSK-sensitive potentiation, we employed two Cre-driver mouse lines for specific hippocampal pyramidal subpopulations, namely G32-4 Cre for CA3 and Amigo2-Cre for CA2 pyramidal neurons. Based on our earlier findings indicating hemispheric asymmetry, we focused on COR. Recording conditions were consistent with previous optogenetic experiments (Fig. 4A, B). To achieve subfield-specific expression, AAV9-hsyn-flex-rc-ChrimsonR-eGFP was injected into either CA3 or CA2 of the right hemisphere (Fig. 4C, D). CA3 COR exhibited notable variability in FSK responses. While the overall mean increase was moderate (110.6 ± 15.6%, *n* = 15, *N* = 5), this reflected again a bimodal distribution: 4 out of 15 recordings (26.7%) surpassed the 20% potentiation threshold (mean: 128.8 ± 4.6%), while the remaining 11 recordings (73.3%) showed minimal to no change (mean: 103.5 ± 1.1%; Fig. 4E). This pronounced variability reflects the heterogeneity of cAMP sensitivity observed in recordings from COR in WT mice, reinforcing the hypothesis that a subpopulation of CA3 neurons is responsible for the cAMP-sensitive component of commissural transmission. In contrast, recordings from CA2 COR revealed minimal cAMP responsiveness. FSK application did not produce potentiation above the 20% threshold in 9 of 10 recordings (mean: 91.7 ± 12.1%). Only one recording (10%) exhibited a substantial increase in synaptic transmission (123.2 ± 15.5%), resulting in a modest overall group mean (94.9 ± 13.7%, *n* = 10, *N* = 4; Fig. 4F). The lack of strong responses across most recordings suggests limited variability and a general insensitivity of CA2-originating commissures to cAMP modulation. Although one CA2-derived recording did show potentiation, statistical comparison of overall responses revealed a significant difference between CA2 and CA3 commissural inputs (*p* = 0.017, unpaired t-test; Fig. 4G), favoring CA3 as the origin of cAMP-sensitive pathways. A bootstrap analysis further supported this distinction, yielding a Cohen’s d of 1.138, indicative of a large effect size and strong functional divergence between these two subfields (Fig. 4I). Together, these findings identify CA3 COR as the likely source of the variable yet pronounced cAMP-dependent potentiation observed at commissural synapses in the left CA1.

**Figure 4.**
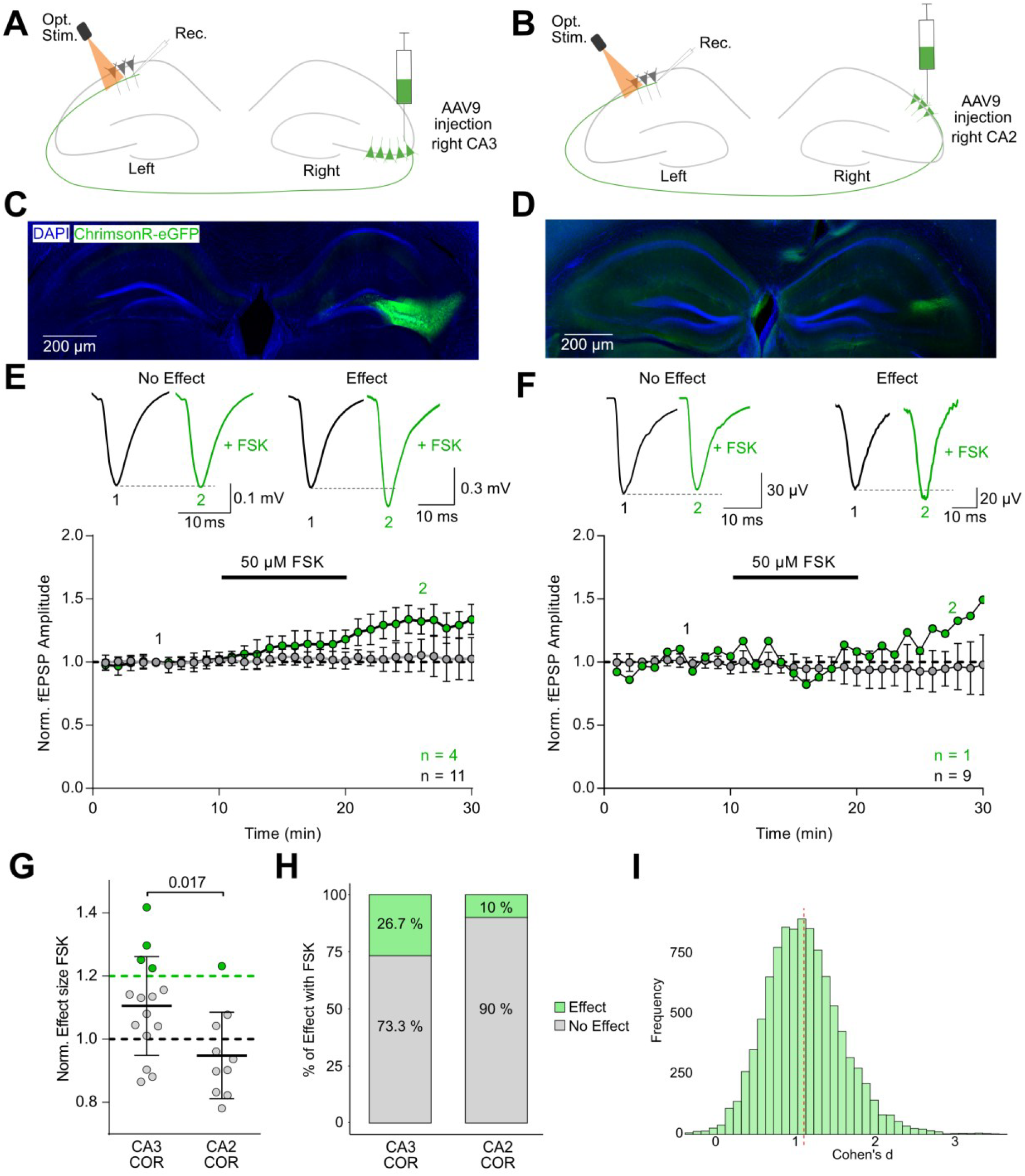
cAMP-induced potentiation is specific to a subset of CA3 commissures originating form the right (COR). **A, B** Schematic of optogenetic stimulation targeting right-hemisphere CA3 **(A)** or CA2 **(B)** pyramidal neurons using G32-4-Cre and Amigo2-Cre mice, respectively. **C, D** Fluorescence images showing targeted ChrimsonR-eGFP expression in CA3 **(C)** and CA2 **(D)** regions, with DAPI counterstaining for cell nuclei. **E, F** Representative optogenetically evoked fEPSP traces evoked by optical stimulation of commissures originating from the right CA3 **(E)** and right CA2 **(F)**. Black traces represent baseline responses; green traces show post-FSK responses. Time course plots from CA3 **(E)** and CA2 **(F)** commissural recordings. Time courses highlight recordings with >20% potentiation (green) versus non-responders (grey), underscoring the higher variability and responsiveness in the CA3 group. **G** Summary of normalized fEPSP amplitudes in response to 50 μM FSK reveals significantly greater potentiation in recordings from CA3-derived commissural projections compared to CA2 (*p* = 0.017, unpaired t-test). **H** Percent stacked bar plot categorizing recordings by response magnitude. In the CA2 group, only 1 of 10 recordings (10%) exceeded the 20% potentiation threshold. In contrast, 4 of 15 CA3 recordings (26.7%) showed pronounced potentiation, highlighting greater variability and a distinct subpopulation of responsive inputs. **I** Bootstrapped distribution of Cohen’s d effect sizes comparing CA3 and CA2 responses indicates a large effect (Cohen’s d = 1.138), supporting a functional distinction favoring CA3 COR.

## Discussion

In this study, we investigated cAMP-dependent plasticity of transmission at glutamatergic synapses in CA1 and found that potentiation was asymmetrically distributed across hemispheres—specifically enhancing transmitter release from commissures originating from the right hippocampus. This lateralization underscores a nuanced role for interhemispheric hippocampal communication in supporting hemisphere-specific contributions to cognition. The specificity and variability of the cAMP-induced potentiation likely reflect the underlying heterogeneity of the hippocampal circuitry. Pyramidal neurons in both CA3 and CA1 are increasingly recognized as heterogeneous populations, structured along multiple spatial axes—transverse (proximal–distal), radial (superficial–deep), and longitudinal (dorsal– ventral). Recent transcriptomic, anatomical, and physiological studies have revealed that subpopulations of CA1 and CA3 neurons differ markedly in gene expression, intrinsic excitability, afferent input, and functional output (Mizuseki *et al*., 2011; Cembrowski, Wang, *et al*., 2016; Sun *et al*., 2017; Cembrowski, 2019). Similarly, CA3 pyramidal cells vary significantly along the proximodistal axis, with CA3b neurons exhibiting strong recurrent excitation and weaker inhibition, favoring synchronous network activity (Sun *et al*., 2017). Morphological and electrophysiological distinctions between thorny and athorny pyramidal neurons further reveal subtypes with differing input sources and roles in sharp-wave ripple initiation (Hunt *et al*., 2018). Such cellular diversity within CA3 likely shapes the input specificity and variability observed in commissural plasticity. Adding a molecular layer of complexity, adenylyl cyclase isoforms—the enzymes responsible for cAMP production—are differentially expressed and regulated across hippocampal neurons (Shahoha *et al*., 2022). This spatial heterogeneity in cAMP synthesis may contribute to preferential activation of cAMP signaling cascades in distinct cell types and subcellular compartments. Even modest presynaptic modulation via cAMP could yield significant functional changes when coupled with postsynaptic reinforcement mechanisms, as suggested by the cooperative model of LTP (Lisman and Raghavachari, 2006), emphasizing the synergistic role of presynaptic neurotransmitter release and postsynaptic AMPA receptor insertion in sustaining plasticity. However, as we consistently used low frequency (0.1 Hz) stimulation for evoking transmitter release, it seems unlikely that postsynaptic depolarization unlocked plasticity-mechanisms that could act synergistically with FSK/cAMP.

The adaptive significance of commissural plasticity is especially compelling in light of its lateralization. While often overshadowed by ipsilateral hippocampal pathways, commissural projections are essential for coordinating activity between hemispheres. The asymmetric nature of the cAMP-mediated potentiation observed here mirrors structural and functional lateralization in the hippocampus. Behavioral studies in rodents have linked paw preference to hemispheric differences in mossy fiber projections and learning strategies (Collins, 1975; Lipp *et al*., 1996). Human split-brain studies revealed that commissural pathways are critical for the integration of memory, perception, and executive function (Gazzaniga, 2005). Genetic models with disrupted forebrain commissures also demonstrate impairments in memory consolidation (Ribeiro, Eales and Biddle, 2013), supporting the role of interhemispheric communication in cognition. Lateralized synaptic plasticity may thus enable functional specialization between hemispheres. For instance, Jordan et al. (Jordan, 2020) propose that asymmetric plasticity could underlie divergent spatial and contextual processing roles in left and right hippocampi. In this context, plasticity at commissural synapses may fine-tune bilateral coordination and enable flexible behavior in response to complex environments.

In summary, our findings demonstrate that commissural projections can engage cAMP-dependent plasticity in a hemisphere-specific manner, likely driven by the interaction of diverse neuronal subpopulations and spatially restricted molecular signaling. This form of plasticity may serve not only as a mechanism for synaptic modulation but also as a substrate for lateralized memory encoding and interhemispheric integration. Future investigations using refined cell-type- and projection-specific manipulations will be essential to unravel how commissural plasticity shapes hippocampal function and contributes to cognition.

## Methods

### Animals

All experiments were conducted in accordance with Directive 2010/63/EU on the protection of animals used for scientific purposes and approved by the Berlin state authorities (LAGeSo; license numbers G0030/20, T0100/03). Mice were bred at the Charité animal facility and housed in individually ventilated cages (4–10 per cage) under a 12-hour light-dark cycle with *ad libitum* access to food and water. Transgenic animals (Amigo2-Cre (RRID:IMSR_JAX:030215) and Grik4-Cre (RRID:IMSR_JAX:006474) mouse lines) were obtained from Jax laboratories and genotyped according to published protocols.

### Stereotactic Viral Injection

Stereotactic injections were performed using a Neurostar half-automated system. Mice received Metamizol (200 mg/kg bodyweight) in drinking water one day before and three days after surgery. Mice were anesthetized with 5% isoflurane, maintained at 3–3.5% throughout surgery. After scalp disinfection and local lidocaine administration, craniotomies were performed. Injection coordinates were: (CA3: AP −1.75 mm, ML 2.1 mm, DV 2.18 mm, CA2: AP −1.75 mm, ML 2.05 mm, DV 1.80 mm). AAV9 encoding Syn-ChrimsonR-tdTomato or Syn-flex-rc-ChrimsonR-GFP (Addgene #84480, a kind gift from Edward Boyden was injected into area CA3 or CA2 via a Hamilton syringe. AAV was injected (200– 400 nL, 40–100 nL/min), followed by a 5-minute wait before needle retraction. Mice received Carprofen (5 mg/kg bodyweight) post-surgery, and NaCl was administered subcutaneously if procedures exceeded one hour. The scalp was sutured, sanitized, and mice were monitored on a heating pad before returning to their home cages.

### Acute Brain Slice Preparation

Acute coronal hippocampal slices (300 µm) were obtained from 6–20-week-old mice from both sexes. Following deep anesthesia with isoflurane, mice were decapitated, and brains were rapidly transferred to ice-cold, oxygenated sucrose-based artificial cerebro spinal fluid (S-aCSF), containing: 124 mM NaCl, 3 mM KCl, 3 mM MgCl_2_, 0.5 mM CaCl_2_, 1.25 mM NaH_2_PO_4_, 26 mM NaHCO_3_, 10 mM glucose, 243 mM sucrose. After a 3-minute incubation, the brain was mounted on a Leica VT1200S vibratome for slicing under continuous ice-cold S-aCSF perfusion. Following dissection, the hemispheres were separated, and a cortical cut was introduced in the left hemisphere to enable reliable identification of laterality. Slices were transferred to a 32°C oxygenated S-aCSF holding chamber for 30 minutes, then allowed to recover in a second chamber with aCSF for ≥1 hour before recordings. ACSF containing: 124 mM NaCl, 3 mM KCl, 1.5 mM MgCl_2_, 2.5 mM CaCl_2_, 1.25 mM NaH_2_PO_4_, 26 mM NaHCO_3_, and 10 mM glucose.

### Electrophysiology

Slices were transferred to a submerged recording chamber perfused with oxygenated aCSF (2–3 mL/min, room temperature). Glass electrodes (1-2 MΩ) filled with aCSF were positioned in CA1 stratum radiatum for field fEPSP recordings. Cells were visualized via IR-DIC microscopy, and recordings were obtained using a MultiClamp 700B amplifier. Schaffer collaterals were stimulated via a bipolar electrode, delivering paired pulses with 40 ms interstimulus interval at 0.1 Hz. Stimulation strength was set to elicit 40–50% of the maximum fEPSP amplitude. Signals were amplified, low-pass filtered (3 kHz), digitized (10 kHz) using a Digidata 1440A, and recorded using pClamp10.

### Optogenetic Stimulation

Photostimulation was achieved using a pE300 CoolLED light source, coupled to an Olympus BX51WI microscope via a liquid light guide and filtered by a HC-Tripleb 365-380-470-585 filter (AHF F66-L422). Synaptic responses were evoked by paired 590 nm light pulses at 0.1 Hz (0.5–2 ms flash duration, 40 ms interval, 1–6 mW/mm^2^, 40× objective (Olympus LUMPL FLN 40XW Objective).

### Data Analysis

Electrophysiological data were analyzed offline using AxoGraph X and GraphPad Prism 6. Baseline normalization, a 1 kHZ low-pass filter, and fEPSP amplitude detection were performed in AxoGraph X. Data normality was assessed using the Shapiro-Wilk test. Statistical analyses were generated in GraphPad Prism 6, with results expressed as mean ± SD. Graphical representations were generated in GraphPad Prism 6 and Affinity Designer 2. To quantify the standardized difference between two independent groups we calculated Cohen’s d and estimated its sampling distribution using nonparametric bootstrapping. The bootstrap procedure involved resampling with replacement from each group’s data to generate 10.000 paired bootstrap samples. For each iteration, Cohen’s d was computed as the difference in means divided by the pooled standard deviation. This process was repeated 10.000 times to construct a distribution of Cohen’s d values. The mean of the bootstrap distribution was taken as the point estimate of the effect size. All analyses and visualizations were conducted using R (version 4.1.1) with the *boot* and *ggplot2* packages.

## List of abbreviations

COL: 
COR: 
fEPSPs: 
FSK: 
Iv: 
LTP: 
PPR: 

## Declarations

### Ethics approval

Not applicable.

### Consent for publication

Not applicable.

### Availability of data and materials

The datasets used and/or analyzed during the current study are available from the corresponding author on request.

### Authors’ contributions

L.F., A.S., and S.O. performed the experiments. L.F., and B.R.R. analyzed the data. B.R.R., D.S. and J.B. conceptualized the research. B.R.R, D.S. acquired funding for the project. B.R.R, D.S. and J.B. supervised the work. L.F. and B.R.R. wrote the paper with input from all authors. All authors approved submission of the manuscript.

## Acknowledgements

The authors thank Anke Schönherr, Susanne Rieckmann, Anny Kretschmer and Katja Czieselsky for excellent technical assistance, and the viral core facility (VCF) of the Charité for providing AAVs.

